# Phenotypic switch and reduced the growth of melanoma spheroids in the presence of mast cell-conditioned medium: potential impact of nutrient starvation effects

**DOI:** 10.1101/2022.09.13.507791

**Authors:** Mirjana Grujic, Thanh Nguyen, Tifaine Héchard, Helen Wang, Maria Lampinen, Aida Paivandy, Gunnar Pejler

## Abstract

Mast cells are abundant in melanoma tumors, and studies suggest that they can be either detrimental or protective for melanoma growth. However, the underlying mechanisms are not fully understood. Here, we adopted a hanging drop-established spheroid system to investigate how mast cells can influence melanoma growth and phenotype in a 3-D context. In the presence of mast cells or mast cell-conditioned medium, melanoma spheroid growth was profoundly reduced. To address the underlying mechanism, we conducted a transcriptomic analysis, which revealed that mast cell-conditioned medium had extensive effects on the melanoma gene expression patterns. Pathway analyses revealed profound effects on the expression of genes related to amino acid and protein metabolism. The conditioned medium also induced an upregulated expression of cancer-related genes, including adhesion molecules implicated in metastatic spreading. In line with this, after transfer to a Matrigel extracellular matrix milieu, spheroids that had been developed in the presence of mast cell-conditioned medium displayed enhanced elevated growth and adhesive properties. However, when assessing for possible effects of nutrient starvation, i.e., reduced nutrient content in mast cell-conditioned medium, we found that the observed effects on melanoma spheroid growth potentially could be explained by such effects. Hence, it cannot be excluded that the observed phenotypic alterations of melanoma spheroids grown in the presence of mast cells or mast cell-conditioned media are, at least partly, due to nutrient starvation rather than to the action of factors secreted by mast cells. Instead, our findings may provide insight into the effects on gene expression events that occur in melanoma tumors under nutrient stress.

## Introduction

Mast cells (MCs) are immune cells with an important role in the innate defense against various external insults such as toxic substances released from venomous animals ^1^, but they are mostly known for their detrimental impact in allergic conditions including asthma ^2^. In addition, MCs are associated with a range of other types of pathological conditions, including arthritis, diabetes, cardiovascular complications and they are also implicated in cancer ^3^. A role of MCs in cancer was originally suggested based on clinical observations of extensive MC infiltration in tumor settings such as breast cancer, lung cancer, pancreatic cancer, Hodgkin’s lymphoma, prostate cancer and malignant melanoma ^4-8^. In these conditions, MCs typically accumulate in large numbers in the tumor stroma, but they can also be found, although usually in lower amounts, in the tumor parenchyma ^4-8^.

In many malignant settings, MC infiltration has been linked to poor clinical outcome, and this has been interpreted as a detrimental impact of MCs under such circumstances. However, there are also numerous studies showing the opposite, i.e., that MC presence associates with good prognosis, suggesting a beneficial role of MCs. Intriguingly though, in some types of malignancies such as melanoma and prostate cancer, there are reports supporting both detrimental and protective functions of MCs ^4-8^.

The role of MCs in cancer has also been addressed by adopting various animal models of MC deficiency and, in many such studies, MCs have been suggested to promote tumor growth ^4, 5^. However, several of these studies were performed in mice in which MC deficiency is accompanied by defects in other cellular niches in addition to their MC-deficiency, and it has thus not been certain that observed effects indeed are due to the MC deficiency vs. effects on other affected cell populations ^9^. However, new generation MC-deficient mice in which only MCs are targeted have been developed ^10^. By evaluating such mice in a lung colonization melanoma model, we showed that the absence of MCs led to reduced lung colonization of melanoma, i.e., supporting a detrimental role of MCs in melanoma dissemination ^11^. However, when assessing mice in which MCs lack the expression of a panel of major granule-localized MC proteases (tryptase, chymase, carboxypeptidase A3), we found that the lung colonization of melanoma was enhanced ^12^. In agreement with the latter findings, we found that mice lacking tryptase displayed enhanced melanoma growth in a subcutaneous tumor model ^13^. Intriguingly, these data collectively suggest that, although MCs overall may support tumor growth, individual compounds expressed by MCs can have a dampening effect on tumor progression.

Altogether, the findings described above suggest a highly complex impact of MCs on tumor cell populations, but the mechanisms underlying these effects have not been fully outlined. Hence, more detailed insight is needed to explain how MCs can affect various aspects of tumor progression. One strategy for gaining such mechanistic understanding can be to implement detailed analyses of the direct interaction between MCs and tumor cells in a milieu reminiscent of the *in vivo* context. Here we developed a spheroid-based 3-D system for this purpose. Our findings reveal that MCs have a dampening effect on melanoma cell growth, which is accompanied by the induction of gene expression patterns that may promote a metastatic phenotype of the melanoma cells.

## Methods

### Bone marrow-derived MCs

Bone marrow-derived MCs (BMMCs) were obtained by culturing bone marrow cells from the femur and tibia of either wild-type, serglycin^-/-^ ^14^ or triple-knockout (Mcpt4^-/-^, Mcpt6^-/-^, Cpa3^-/-^) ^15^ mice (all on C57BL/6J genetic background) as previously described ^16^ but with the following modifications: cells were cultured in RPMI-1640 medium supplemented with 10% heat-inactivated FBS, 0.1 mM nonessential amino acids, 50 μM b-mercaptoethanol, 1 mM sodium pyruvate, 10 mM HEPES, 2 mM L-glutamine, 100 U/ml penicillin, 100 μg/ml streptomycin, 20 ng/ml recombinant IL-3 and 20 ng/ml stem cell factor (Peprotech Nordic, Sweden).

### Preparation of conditioned medium from human skin mast cells

Human MCs were isolated from healthy skin as described previously ^17^, and cultured for five days in RPMI (Gibco/Thermo Fisher Scientific, Waltham, MA) supplemented with 10% heat-inactivated FCS, 4 mM L-glutamine, SCF (100ng/mL), IL-4 (20ng/mL) and antibiotics. After incubation, cells were pelleted by centrifugation and removed from the medium.

### Development of spheroids

The mouse melanoma cell line B16.F10 (ATCC; CRL-6475) was a gift from A.R. Thomsen (Copenhagen University, Denmark) and the human melanoma cell line MM253 was purchased from CellBank Australia (CBA; CBA-1347). B16.F10 and MM253 cells were cultured in DMEM or RPMI medium, respectively, both supplemented with 10% FBS (cat. #10500064, Thermo Scientific, Waltham, MA), 2 mM Glutamine (cat. #G7513-100ml, Merck KGaA, Darmstadt, Germany) and 1% penicillin and streptomycin solution (cat. #P0781-100ml, Merck KGaA). At 60-90% confluency, the cells were trypsinized. B16.F10 cells were then resuspended in RPMI medium with 20 ng/ml rm IL-3 and 20 ng/ml rm SCF, whereas MM253 were resuspended in RPMI medium with 20 ng/ml rh IL-4 and 100 ng/ml rh SCF, and counted using trypan blue in order to adjust the cell concentration to 80,000 cells/ml. The cell suspensions were supplemented with 0,25% methyl cellulose (cat. #M7027, Merck KGaA), and with either BMMCs in RPMI medium (with 20 ng/ml rm IL-3 and 20 ng/ml rm SCF) or with conditioned medium from BMMCs, or with conditioned medium from human skin MCs. 25 µl of the prepared cell suspensions, corresponding to 2000 B16.F10/MM253 cells, were pipetted on a Petri dish lid, which was then inverted onto a Petri dish filled with sterile PBS. Petri dishes with hanging drops were placed in a CO_2_ incubator for 3 or 5 days. At chosen time points, pictures of spheroids in hanging drops were taken with a Nikon TMS-F Inverted Phase Contrast No.300630 microscope (Nikon, Tokyo, Japan), using 2x objective and a Nikon D3300 camera. Spheroids were transferred into 1,5 ml centrifuge tubes; 100 µl of trypsin-EDTA solution (Merck KGaA; cat. #T3924) was added and the samples were trypsinized on a bench top shaking incubator (Corning LSE, NY) at 37°C, 190 rpm, for 1h. Before counting the cells with a hemocytometer, the spheroids were additionally disrupted by pipetting and 10 µl of trypan blue (cat. #15250-061, Gibco, Paisley, UK) was added to the samples. When the cells were subjected to the flow cytometry analysis, 1 ml of RPMI medium with 10% FCS was added to the samples after 1h trypsinization.

### RNA extraction, quantitative RT-PCR, ddPCR analysis

RNeasy Micro Kit (Qiagen GmbH, Hilden, Germany; cat. #74004) was used to isolate total RNA from the spheroids (containing from 5,000 to 20,000 cells). Total RNA concentration and purity were measured using a NanoDrop 1000 Spectrophotometer (Thermo Scientific) and the ND-1000 V3.7.0 program. cDNA synthesis and qPCR analysis was performed as described ^13^. Gene expression levels were presented relative to the house keeping gene (glyceraldehyde 3-phosphate dehydrogenase; GAPDH) and relative to nontreated B16.F10 spheroids. ddPCR analysis was performed with EvaGreen Supermix (Bio-Rad; cat. #1864033) and the Bio-Rad ddPCR QX200 system (Bio-Rad) according to the manufacturer’s instructions. Briefly, 12.5 µl of Eva-Green master mix (Bio-Rad), 1 µl HindIII-HF (NEB, Hitchin, UK), 10.5 µl of cDNA (5ng/µl per reaction), and 1 µl of respective primer pairs (200 nM final concentration) were mixed and distributed into 96-well qPCR plate (Bio-Rad; cat. #12001925). Droplets were generated in an Automated Droplet Generator (Bio-Rad). PCR was performed with a hot-start/enzyme activation at 95°C for 5 min, denaturation at 94°C for 30s, and amplification at 58°C for 1 min over 40 cycles, followed by signal stabilization at 4°C for 10min and 90°C for 5 min. For all steps, a ramp rate of 2°C/s was used. The droplets were analyzed in a QX200 droplet reader (Bio-Rad) and the data were analyzed with QuantaSoft Analysis Pro1.0.569 (Bio-Rad). Results were presented as number of copies of a transcript of interest per µl.

The following primer pairs were used: GAPDH, forward primer (5’-3’): CTC CCA CTC TTC CAC CTT CG, reverse primer (5’-3’): CCA CCA CCC TGT TGC TGT AG; DCT, forward primer (5’-3’): TCC TCC ACT CTT TTA CAG ACG, reverse primer (5’-3’): ATT CGG TTG TGA CCA ATG GG; GP100, forward primer (5’-3’): AGC ACC TGG AAC CAC ATC TA, reverse primer (5’-3’): GTT CCA GAG GGC TGT GTA GT; β-actin, forward primer (5’-3’): AGA CAG CAC TGT GTT GGC ATA GAG, reverse primer (5’-3’): AGG TCA TCA CTA TCG GCA ATG AGC.

### Transcriptome analysis

Transcriptome analysis was performed using the Ion AmpliSeq™ Transcriptome Mouse Gene Expression Kit (Life Technologies, Carlsbad, CA), followed by analysis as described ^18^. R (version 4.1.2) was used.

### Flow cytometry analysis

After trypsinization, cells were rinsed with a cold FACS buffer (PBS supplemented with 2% FCS). For apoptosis analysis, cells pooled from several spheroids were recovered in a cold binding buffer (BD Biosciences, Franklin Lakes, NJ; cat. #556454) and stained with 3 µl of FITC-Annexin V (BD Biosciences; cat. #556419) and 3 µl Draq7 (Biostatus, Shepshed Leicestershire, UK: cat. #DR71000). Samples were incubated at RT, in the dark for 15 min. Subsequently, cells were fixed in 1% PFA in FACS buffer at 4°C in the dark and for 15 min. Cells were rinsed with FACS buffer, recovered in FACS buffer and filtered through 70 µm pluristrainer (PluriSelect Life Science: Leipzig, Germany: cat. #43-10070-40) before being acquired by a BD Accuri C6 Plus flow cytometer (BD Biosciences). Data from 15,000 events/sample were analyzed by FlowJo software (Ashland, OR). For cell cycle analysis, cells pooled from several spheroids were fixed in 1% PFA in FACS buffer (as described above), rinsed with FACS buffer and permeabilized in 0.5% saponin (Merck KGaA) in PBS, at RT and for 10 min. Subsequently the cells were stained with Draq7 in 0.5% Saponin in PBS, at RT in the dark and for 15 min. The cells were rinsed with 0.5% Saponin in PBS, recovered in FACS buffer and filtered through 70 µm pluristrainer (PluriSelect Life Science; cat. #43-10070-40). The samples were acquired and analyzed as described above.

### Immunostaining of spheroids and confocal analysis

Hanging drops with formed melanoma spheroids were transferred into 8-chamber culture slides (cat. #C8-1.5H-N, Cellvis, Sunnyvale, CA). The spheroids were covered with 100 µl Matrigel (cat. #356230, Corning, Bedford, MA) and allowed to solidify in a CO_2_ incubator at 37°C for 40 min. Samples were fixed with 300 µl 4% PFA (cat. #sc-281692, Santa Cruz Biotechnology, Dallas, TX) in PBS for 20 min at RT, washed once with IF buffer (PBS with 0.2% Triton and 0.05% Tween 20) and permeabilized with a permeabilization solution (PBS with 0.5% TritonX-100) for 20 min at RT. Samples were rinsed with IF buffer for 5 min and blocked with a blocking solution (1% BSA in IF buffer) for 30 min at RT. The primary antibody rabbit anti-Ki67-Alexa647 (cat. #NB110-89717AF647, Novus Biological, Littleton, Colorado) was diluted in the blocking solution (1:250) and 300 µl of the antibody solution was applied to spheroids in Matrigel. The staining was performed overnight at 4°C in dark. The next day, samples were rinsed three times with IF buffer (5 min/rinsing), counter-stained with NucBlue (cat. #R37605, Invitrogen, Eugene, OR) according to manufactures instructions for 45 min at RT, rinsed three times with IF (5 min/rinsing) and covered with IF. The chambers were kept at 4°C until analysis with a laser-scanning microscope equipped with ZEN 2009 software (LSM700, Carl Zeiss, Berlin, Germany). The area of nucleus (NucBlue+) was determined with Image J software, as well as the intensity of Ki67-staining of the nucleus area. The result was presented as the whole intensity of Ki67-staining divided with the whole nuclear area.

### Growth of spheroids in Matrigel

Spheroids were transferred from hanging drops into with Matrigel precoated 8-chamber culture slides covered with 100 µl Matrigel, and was left to solidify in a CO_2_ incubator at 37°C for 40 min. Samples were covered with 500 µl of complete RPMI medium (supplemented with 10% FCS, 2 mM Glutamine and 1% penicillin and streptomycin solution, 50µM βME (cat. #31350-010, Gibco), 1% sodium pyruvate (cat. #S8636-100ml, Merck KGaA) 1% MEM (cat. #M7145-100ml, Merck KGaA), 1% Hepes (cat. #H3662-100ml, Merck KGaA). Medium was changed every second day. Pictures of spheroids were taken with a Nikon D3300 camera connected to a Nikon TMS-F Inverted Phase Contrast No.300630 microscope, using 2x objective. The area of spheroids was measured with Image J software and the data was presented as % of growth, obtained with the following equation: (((area on day X)-(area on day 0))/(area on day 0))*100.

## Results and Discussion

### MCs impair the growth of melanoma spheroids

To create a 3-D environment reminiscent of the *in vivo* milieu found in the parenchyma of melanoma tumors, B16F10 mouse melanoma cells were cultured in a hanging drop system. Under these conditions, the melanoma cells cluster into spheroids (Fig. 1A), with features that partly can replicate properties of the tumors found *in vivo* ^19, 20^. To assess the impact of MCs, spheroids were developed either in the absence or presence of bone marrow-derived MCs, followed by an assessment of spheroid growth. As seen in Fig. A-B, when MCs were present at a 1:1 ratio to the melanoma cells, spheroid growth was profoundly reduced, hence indicating that MCs have the ability to interfere with melanoma expansion. In agreement with this, measurements of total RNA content revealed that the presence of MCs led to a trend of reduction of RNA content of the melanoma cells (Fig. 1C). Next, we asked whether the reducing impact of MCs on melanoma growth was associated with a reduction in the expression of the melanoma-specific genes: L-dopachrome tautomerase (DCT) and GP100. However, the expression of DCT and GP100 normalized to the expression of GAPDH as house-keeping gene (and related to the expression in nontreated spheroids) was not affected by MC presence (Fig. 1D). When assessing absolute gene expression by using ddPCR, we found that MCs caused a substantial reduction in the expression of both DCT and GP100. However, the expression of house-keeping genes (GAPDH and β-actin) was suppressed to a similar extent. Altogether, these data indicate that MCs have the capacity to reduce melanoma spheroid growth and to cause an overall reduction in gene expression in the melanoma cells.

**Fig. 1.**
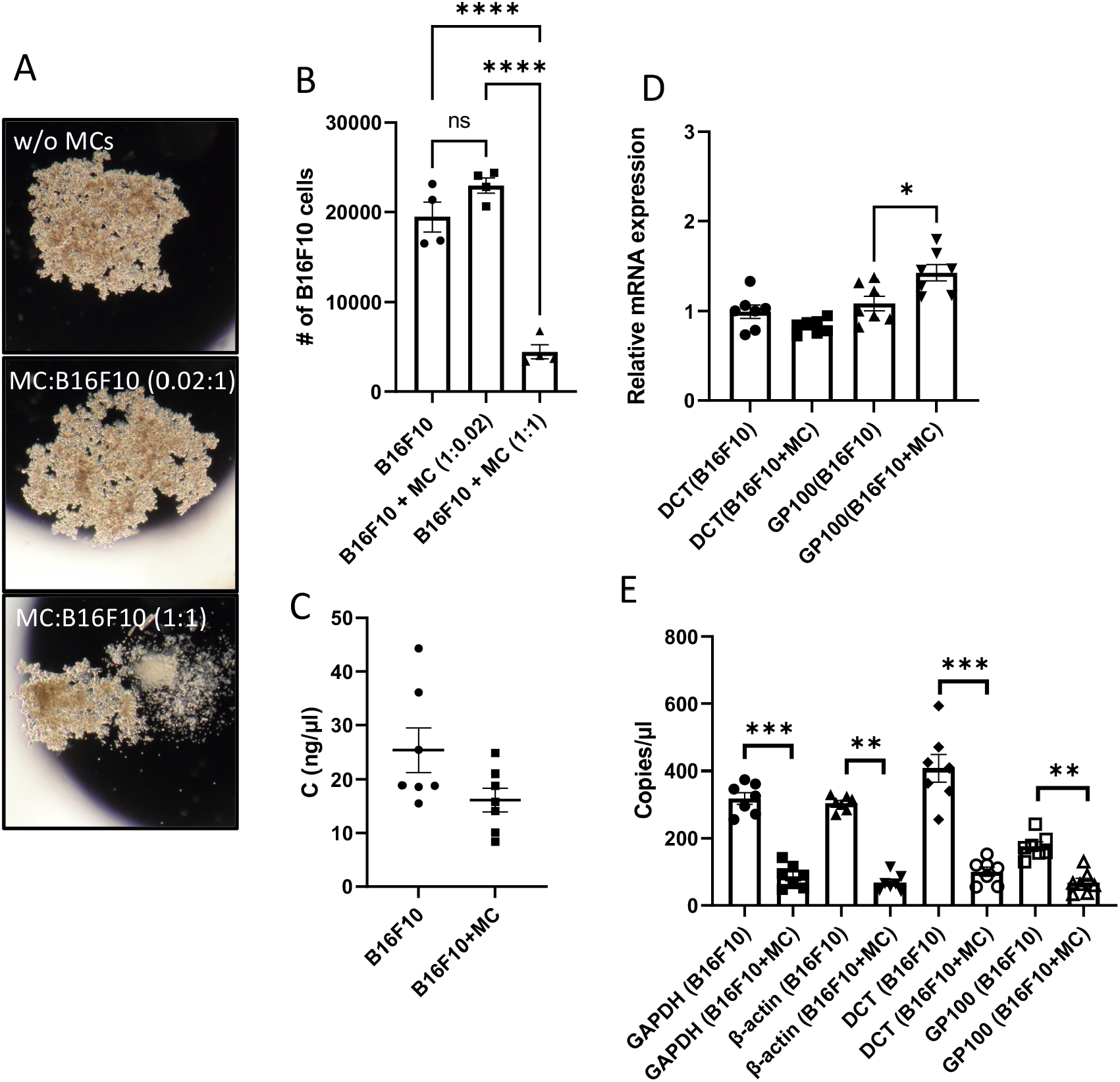
MCs inhibit melanoma spheroid growth. Melanoma spheroids were grown for 5 days in the absence or presence MCs. (A) Images of melanoma spheroids formed in hanging drops were taken with inverted phase contrast microscope. (B) Melanoma cell numbers determined after trypsinization of the spheroids, staining with trypan blue and counting the cells using a hemocytometer. Data are representative of two independent experiments and are given as mean values ± *SEM* (*n* = 4). One-way ANOVA, Tukey’s multiple comparisons test. (C-E) Melanoma spheroids, nontreated or treated with MCs (1:1 ratio of B16F10 and MCs), were harvested and total RNA was isolated. (C) Total RNA concentration. Data shown represent means ± SEM of two independent experiments (n=7). (D) qPCR analysis for the expression of melanoma-specific genes: DCT and GP100. Expression of genes was evaluated relative to GAPDH, and normalized to nontreated spheroids. Results are presented as mean values ± SEM (n = 7) of two independent experiments. Mann-Whitney test. (E) Expression levels of housekeeping genes (GAPDH and β-actin) and melanoma specific genes (DCT and GP100) were measured by ddPCR. Results are presented as mean ± SEM (n = 7) of two independent experiments. Mann–Whitney test. \**p* ≤ 0.05, \*\**p* < 0.01, \*\*\**p* < 0.001, \*\*\*\**p* < 0.0001.

### Reduced growth of mouse and human melanoma spheroids in the presence of MC-conditioned media

The data above suggest that MCs can have a negative impact on melanoma spheroid growth, and this could potentially require MC-melanoma cell-cell contact or be independent on direct contact between the two cell populations. To approach this issue, we assessed if factors present in the MC-conditioned medium can influence melanoma spheroid growth. Indeed, when melanoma spheroids were developed in the presence of normal culture medium with the addition of MC-conditioned media (10-50%), reduced melanoma growth was observed (Fig. 2A-B). To address whether such a dampening effect of MC-conditioned media on melanoma growth also can be recapitulated in a human system, we cultured human melanoma cells (MM253) in the presence of conditioned medium from primary human skin MCs. As seen in Fig. 2C-D, also human melanoma spheroid growth was reduced in the presence of MC-conditioned medium. Altogether, these findings suggest that MCs secrete compounds that have the ability to interfere with the growth of melanoma spheroids.

**Fig. 2.**
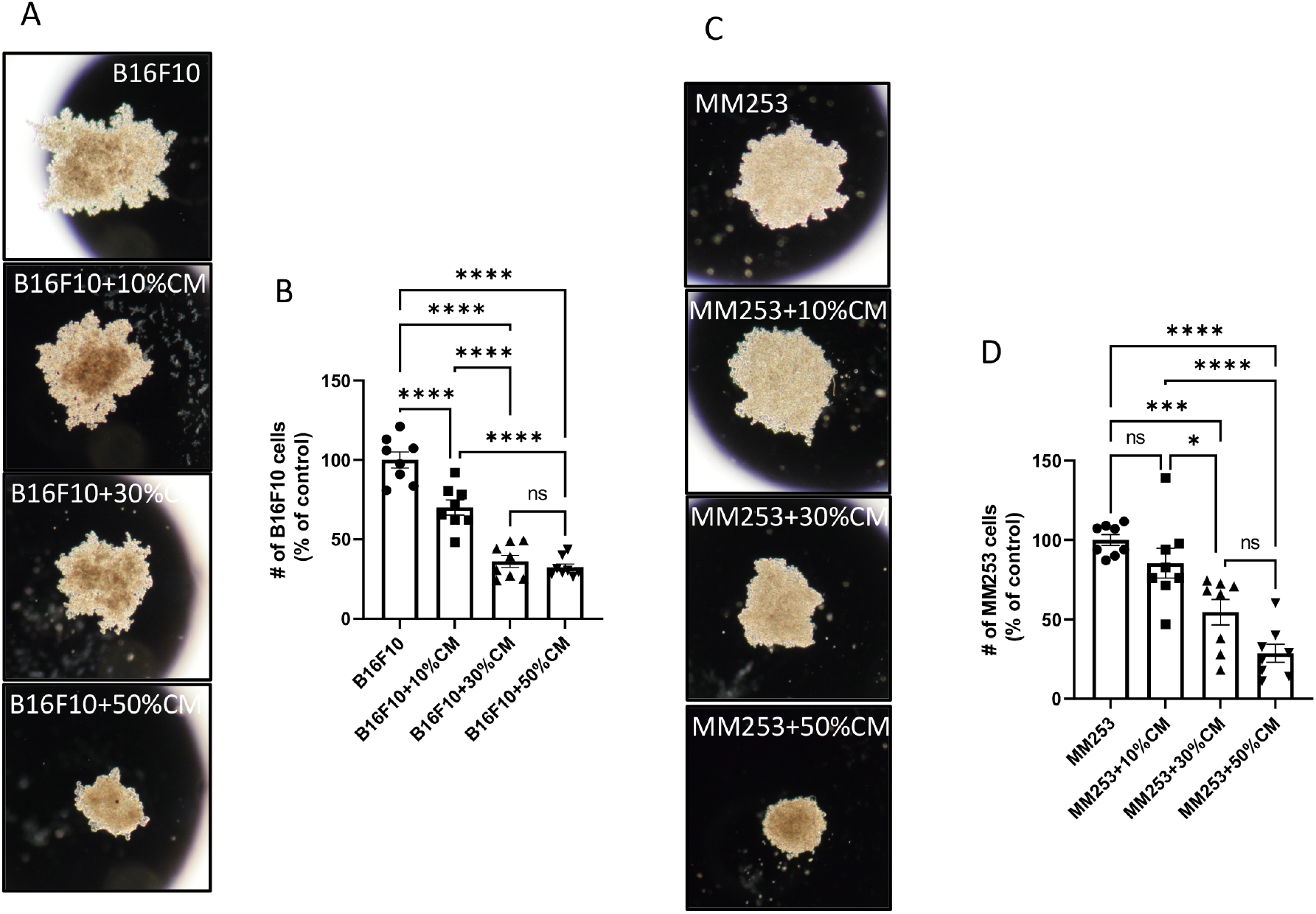
Reduced growth of mouse and human melanoma spheroids in the presence of MC-conditioned medium. Mouse (A, B) or human (C, D) melanoma spheroids were developed for 5 days in the absence or presence of different amounts (10-50%) of MC (BMMCs or human skin MCs)-conditioned media (CM). (B, D) Melanoma cells numbers determined after trypsinization of the spheroids and staining with trypan blue. Data shown represent means ± SEM of two independent experiments. One-way ANOVA, Tukey’s multiple comparisons test. \**P* < 0.05, \*\*\**P* < 0.001, \*\*\*\**p* < 0.0001.

### The negative impact of MCs on melanoma spheroid growth is independent of the MC-restricted proteases and serglycin

MCs are highly granulated cells, with various MC-restricted proteases and serglycin proteoglycans accounting for a major fraction of the total granule content ^21^. To assess whether any of these compounds can influence melanoma growth, we developed melanoma spheroids in the presence of wild-type (WT) MCs or MCs with either multiple deficiency in the MC-restricted proteases (chymase, tryptase, CPA3, ^15^) or deficient in serglycin ^14^. However, the impact of MCs on melanoma growth was not affected by the deficiency in either of these compounds, suggesting that the ability of MCs to impair melanoma spheroid growth is independent of the MC-restricted proteases and on serglycin proteoglycan.

### MC-conditioned medium affects the proliferation of melanoma cells

To provide insight into the mechanism by which MCs affect melanoma spheroid growth, we assessed the effect of MC-conditioned media on the cell cycle regulation in melanoma cells recovered from the spheroids. As seen in Fig. 4A, a substantial fraction of the melanoma cells in non-treated spheroids were in the G2 phase, i.e., in a proliferating state. However, the fraction of cells in G2 phase was significantly reduced in the presence of MC-conditioned media, suggesting reduced proliferation (Fig. 4A), and this was accompanied by a corresponding increase in the population of cells in G1/G0 phase. In agreement with this, a reduction of the number of Ki67-positive cells was seen in melanoma spheroids that had been treated with MC-conditioned medium and then transferred to an extracellular matrix (ECM; Matrigel) context (to preserve spheroid integrity) (Fig. 4B). Potentially, an increased rate of apoptosis could contribute to the observed reduction of melanoma spheroid growth in the presence of MCs or MC-conditioned media. To address this possibility, we additionally stained melanoma cells recovered from the spheroids with annexin V and Draq7. This revealed that MC-conditioned medium evoked a modest decrease in the fraction of viable cells, accompanied by a marginal increase in double annexin V/Draq7-positve (necrotic/late apoptotic) cells (Fig. 4C), suggesting that the MC-conditioned medium caused minimal cell death of the melanoma cells.

**Fig. 3.**
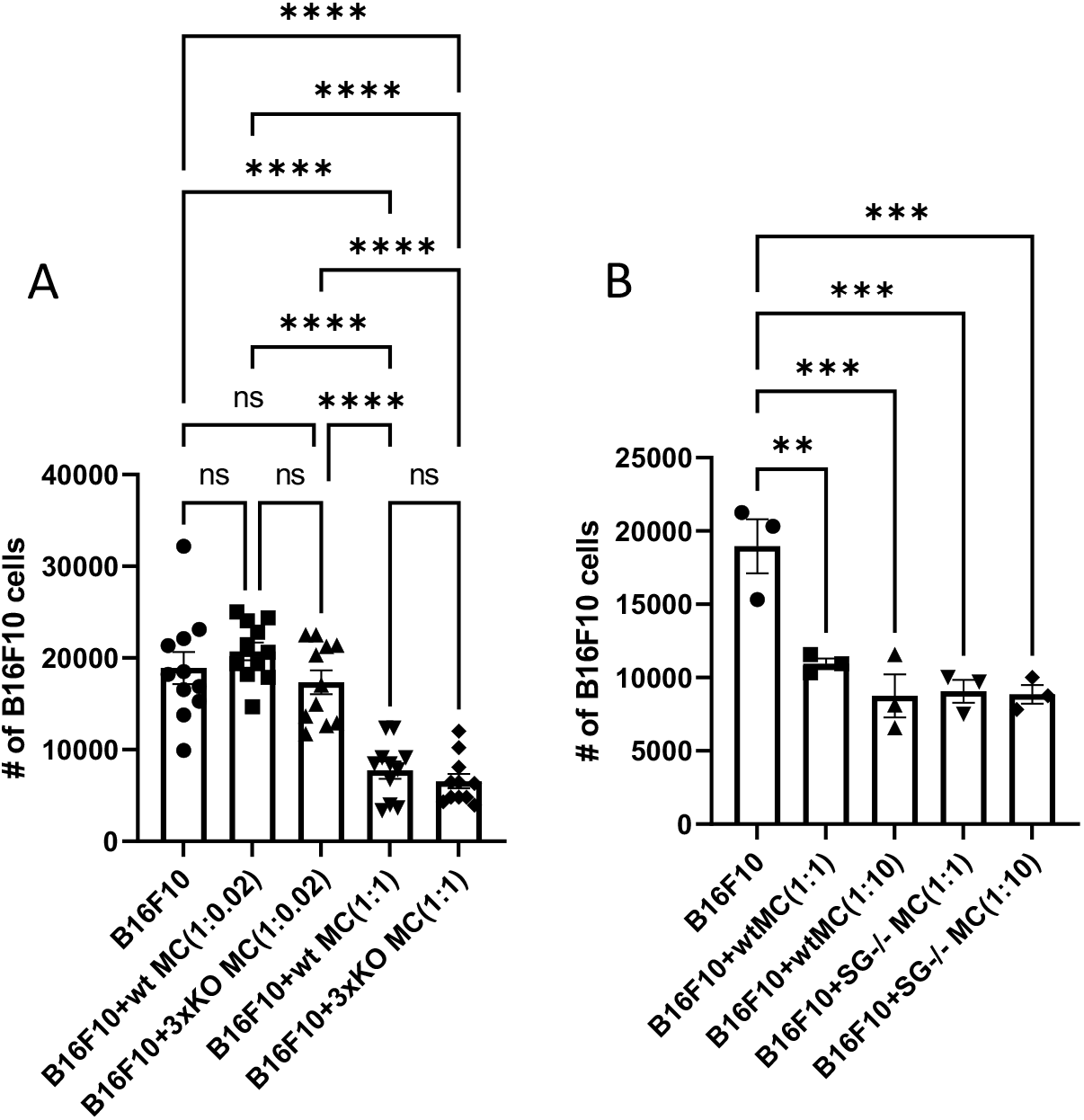
The suppressing effects of MCs on melanoma spheroid growth is independent of serglycin and of major MC proteases. Melanoma spheroids were developed in the absence of MCs, or in the presence of either WT, serglycin^-/-^ or triple knockout (Mcpt4^-/-^; Mcpt6^-/-^; Cpa3^-/-^) MCs. On day 5, melanoma spheroids were harvested, trypsinized, stained with trypan blue and counted with a hemocytometer. Data shown represent means ± SEM (n=11) of three independent experiments (A), or representative of two performed experiments (n=3) (B). One-way ANOVA, Tukey’s multiple comparisons test. \*\**P* < 0.01, \*\*\**P* < 0.001, \*\*\*\**p* < 0.0001.

**Fig. 4.**
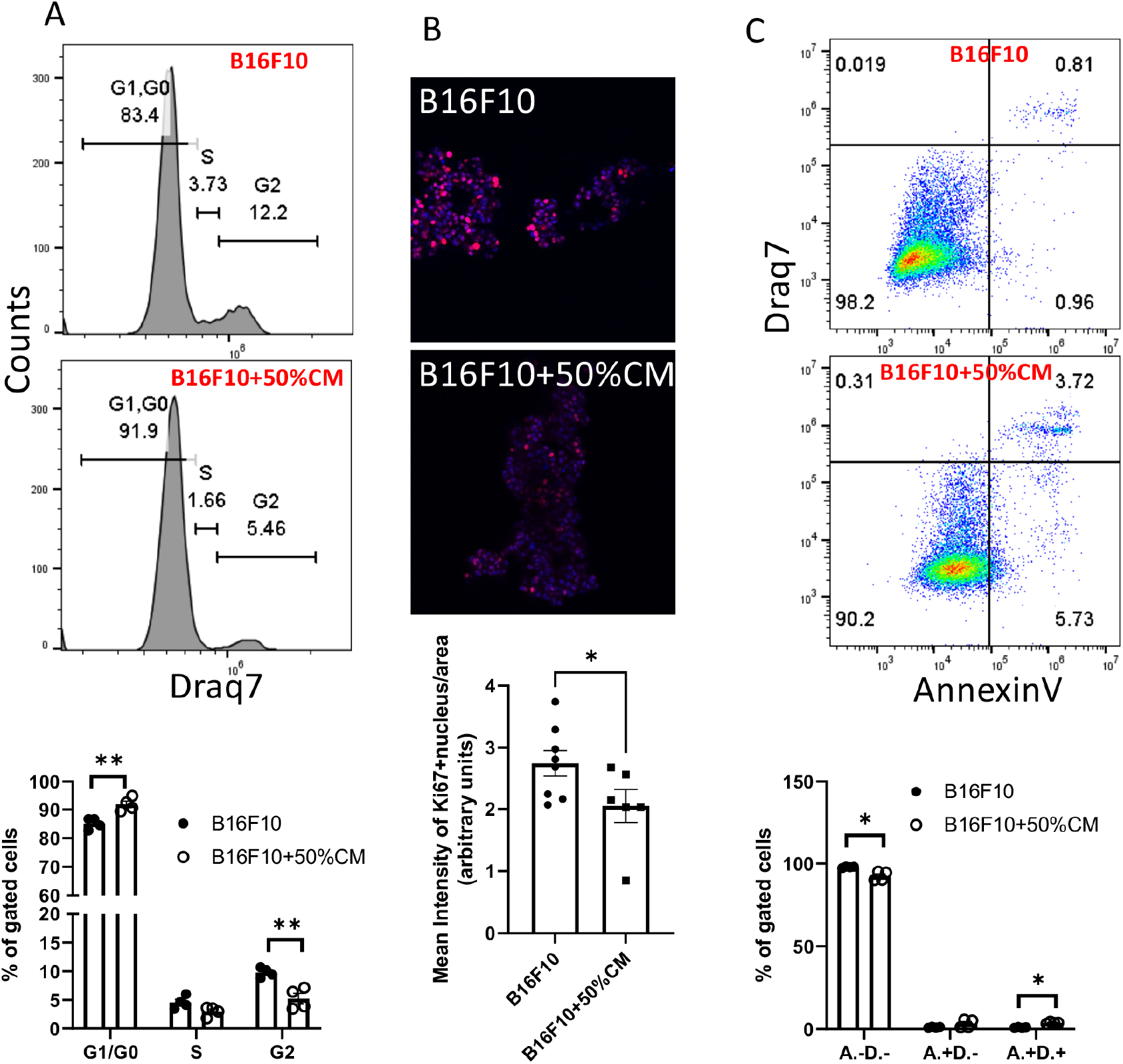
MC-conditioned medium affects the proliferation of melanoma cells. (A-C) Melanoma spheroids were developed in the absence or presence of 50% of MC-conditioned media (CM). On day 3, melanoma spheroids were harvested and either: (B) transferred to Matrigel prior immunostaining for Ki67 and analysis with a confocal microscope; (A,C) trypsinized to a single cell suspension and either (A) permeabilized, stained with Draq7 and analyzed with a flow cytometer or (C) stained with Annexin V (A.) and Draq7 (D.). Shown data are representative of two independent experiments and are given as means ± SEM (n=4-7). Mann-Whitney test, \**P* < 0.05, \*\**P* < 0.01.

### MC-conditioned medium affects the transcriptome of melanoma spheroids

To provide an increased understanding of how MCs can affect melanoma, we next performed a transcriptome analysis to address the effects of MC-conditioned medium on relative gene expression in the melanoma spheroids. For this, melanoma spheroids were grown either in the absence or presence of MC-conditioned media for periods up to 5 days, followed by Ampliseq transcriptome analysis. As seen in a multidimensional scaling plot analysis, the transcriptomes of control spheroids and spheroids treated with MC-conditioned media clustered closely together after 24 h of incubation, indicating minimal effects of MCs on the gene expression patterns at this time point. In contrast, control- and treated melanoma spheroids showed a clear separation after prolonged culture periods (5 days), indicating profound effects of the MC-conditioned media on gene expression patterns (Fig. 5A). These findings were substantiated by a volcano plot analysis, which confirmed both the minimal effects of MC-conditioned medium on gene expression patterns after short term culture, and also the extensive effects seen after longer periods of treatment. As visualized in the volcano plot (Fig. 5C-D), MC-conditioned medium caused strongly downregulated relative expression of numerous genes, but also a profoundly upregulated expression of others. Altogether, 401 genes were significantly downregulated and 533 genes were significantly upregulated in response to the MC-conditioned medium (Suppl Table 1). Effects on gene expression patterns were also visualized in a heat map analysis (Fig. 5B). KEGG pathway analysis of the gene expression patterns revealed that MC-conditioned medium induced an upregulated expression of cancer-related genes, but also of genes with a role in virus (measles) defense. An upregulated expression of genes related to virus defense was also supported by GO Biological Processes analysis, whereas GO Molecular Function pathway analysis revealed an upregulated expression of genes involved in protein binding and transferase activity. Among the pathways that were downregulated by MC-conditioned medium, we noted a preponderance of pathways related to protein and amino acid metabolism, based on both KEGG, GO Biological Process and GO Molecular Function analyses (Fig. 5E-G).

**Fig. 5.**
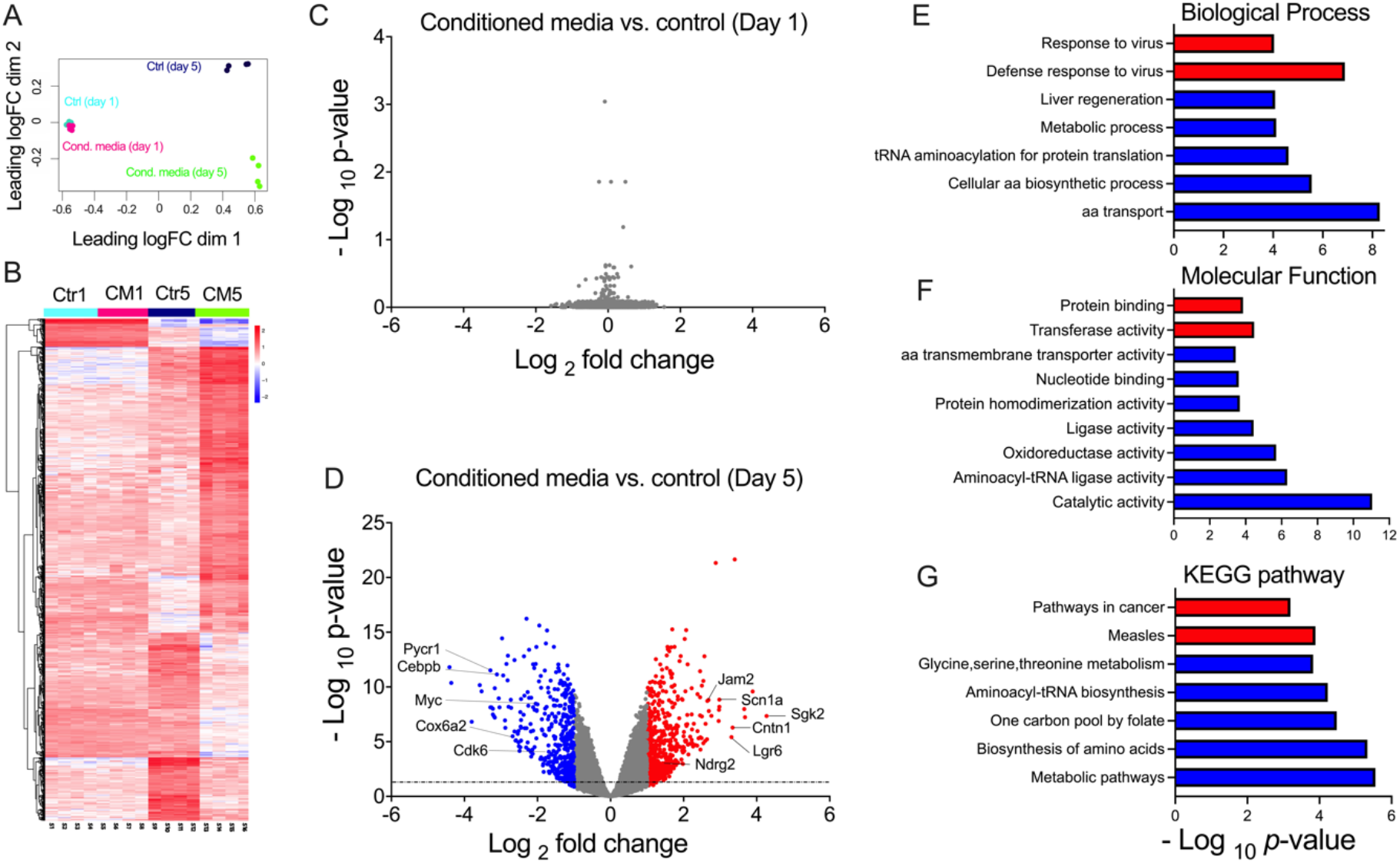
Effects of MC-conditioned medium on transcriptome of melanoma spheroids. Melanoma spheroids were cultured either alone or in the presence of MC-conditioned medium, for 1 or 5 days as indicated, followed by Ampliseq transcriptome analysis. (A) Multidimensional scaling plot displaying the clustering of samples. (B) Heat map displaying hierarchical clustering of differentially expressed genes (DEGs; red represents upregulation and blue downregulation). Original data were scaled to have 0 mean. (C-D) Volcano plots of log2 fold changes of gene expression after treatment of melanoma spheroids with MC-conditioned medium. DEGs with P <0.05 and absolute FC ≥ 1 are indicated in red and DEGs with P <0.05 and absolute FC ≤1 are indicated in blue. The horizontal dashed lines denote P value cutoff of 0.05. (E, F, G) Gene Ontology (GO; http://geneontology.org/) and Kyoto Encyclopedia of Genes and Genomes (KEGG; https://www.genome.jp/kegg/) pathway enrichment analyses for DEGs in melanoma spheroids treated MC-conditioned medium versus untreated spheroids. The GO and KEGG pathway enrichment analyses indicate the P values in all categories. All P values were adjusted using the Benjamini-Hochberg procedure (P < 0.05). aa; amino acids, CM; conditioned medium, Ctr; control.

### Melanoma spheroids developed in the presence of MC-conditioned medium display increased growth and increased adhesive properties after transfer to an extracellular matrix milieu

Fibroblasts are main producers of ECM, and since fibroblasts were not present in our spheroid system, our cultures will have a relatively low content ECM (see also ^22-24^. Hence, in this sense, spheroid systems may mimic the milieu prevailing in the tumor parenchyma. In contrast, the tumor stroma has a substantial content of fibroblasts and is thereby rich in ECM ^25, 26^, and we next asked whether factors released by MCs can influence the properties of melanoma spheroids in an ECM-rich environment. To address this, we transferred control spheroids and spheroids that had developed in the presence of MC-conditioned medium to an ECM milieu (Matrigel). Potentially, this could replicate a setting in which melanoma cells egress from the tumor parenchyma to the stroma, e.g., in the process of metastatic dissemination ^26^. As seen in Fig. 6A, control spheroids were extensively dispersed after their transfer to the Matrigel context. In contrast, spheroids that had been developed in the presence of MC-conditioned medium clumped tightly together after transfer to Matrigel. To monitor the growth capacities of the control spheroids and spheroids developed in the presence of MC-conditioned medium, the expansion of individual melanoma entities was followed over time. As seen in Fig. 6B, melanoma entities derived from spheroids developed in the presence of MC-conditioned medium showed a substantially higher growth capacity in Matrigel, vs. spheroids that were developed in the absence of MC-conditioned medium. Moreover, even after prolonged times of culture in Matrigel, spheroids that were originally developed in the presence of MC-conditioned medium clumped much more tightly together in comparison with spheroids developed in the absence of MC-conditioned medium, compatible with a higher expression of adhesion molecules.

**Fig. 6.**
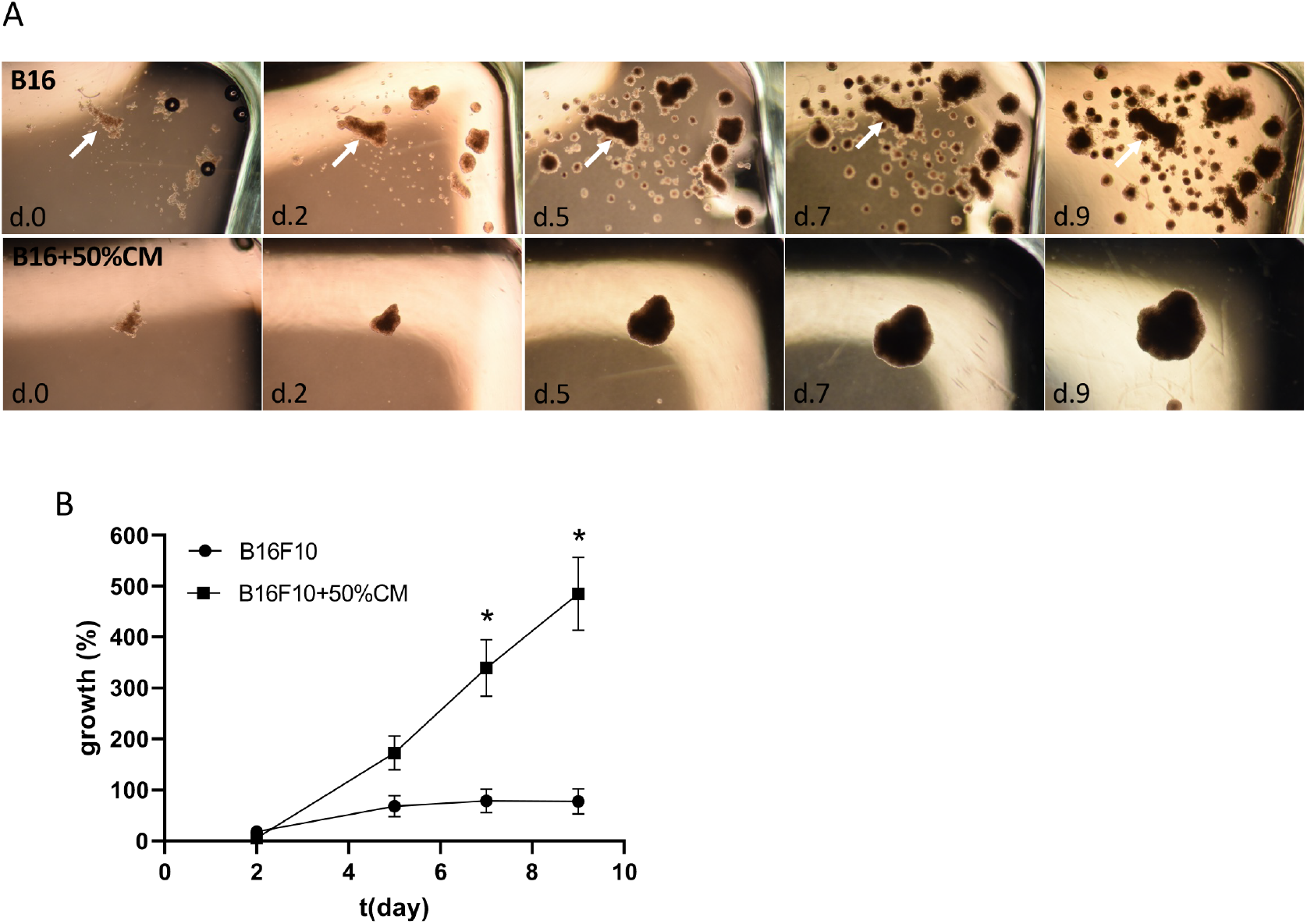
Enhanced growth in Matrigel of melanoma spheroids developed in the presence of MC-conditioned medium. (A) Melanoma spheroids were developed in the absence or presence of 50% of MC-conditioned media (CM). After 5 days, spheroids were transferred into Matrigel, covered with medium, and cultured further for 9 days. Growth of individual spheroids was followed over time. Images were taken at the indicated time points, and spheroid size was recorded. Representative spheroids were used for quantification of growth (arrows in (A)). (B) depicts quantification of the data. Data are representative of three independent experiments and are given as means ± SEM (n=4). Multiple Mann-Whitney tests, \**P* < 0.05.

### The role of nutrient starvation effects in the regulation of melanoma spheroid growth in the presence of MC-conditioned medium

Our findings suggest that MC-conditioned medium affects the growth of melanoma spheroids, implying that MCs secrete factors that affect the melanoma cells. However, although such effects were seen even at low concentrations of MC-conditioned medium (down to 10% of the growth medium; see Fig. 2B), we could not completely exclude that our findings could be attributed to the nutrient consumption that might have occurred within the medium that had been conditioned by MCs. To assess this possibility, we replaced the MC-conditioned media with equal amounts of nutrient-poor buffer (PBS). When assessing the effects of medium supplemented with PBS at various ratios on the melanoma growth, we found that the impairment of melanoma spheroid growth was similar to the effects of MC-conditioned medium (Fig. 7). Hence, these findings do not a support a role of factors secreted by MCs in the regulation of melanoma spheroid growth under the conditions employed in this study. Further, we cannot exclude that the effects of MC-conditioned medium on gene expression patterns in the melanoma spheroids are due to nutrient starvation effects rather than to the presence of factors secreted by MCs. Instead, the noted effects on melanoma phenotype may be attributed to starvation effects. Notably though, nutrient starvation is a common feature of solid tumors ^27^, and this study may thus provide insight into the phenotypic switch that may occur in melanoma tumors under nutrient stress.

**Fig. 7.**
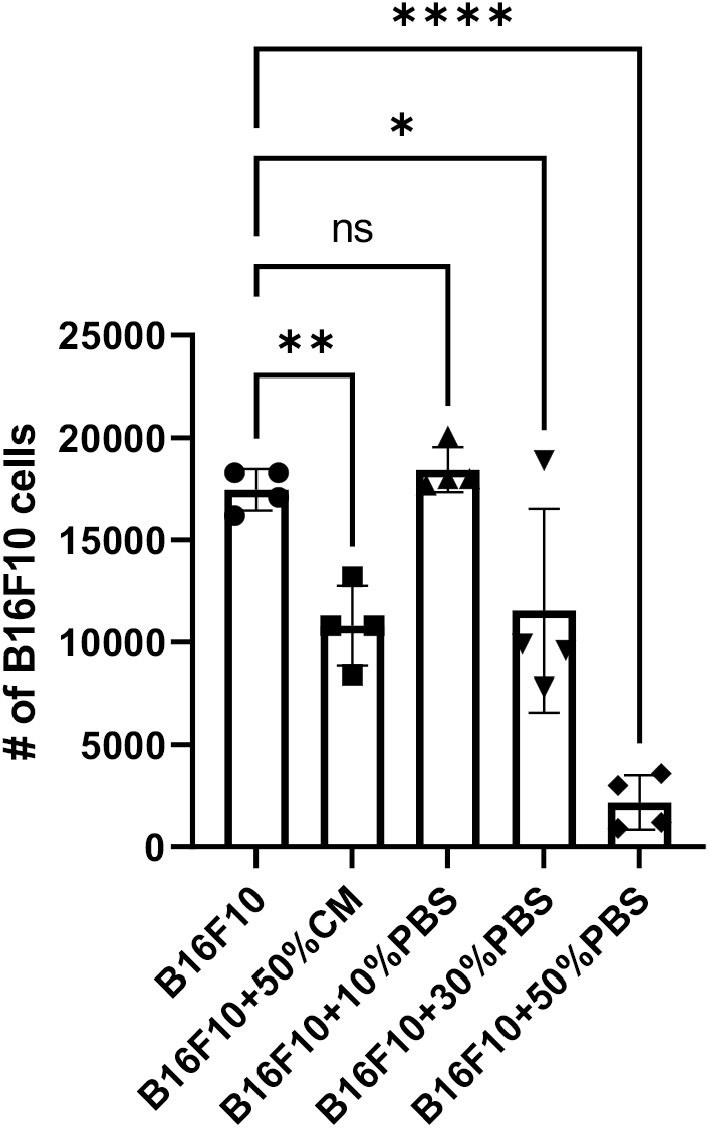
Nutrient starvation has similar suppressing effect on melanoma spheroid growth as MC-conditioned medium. Mouse melanoma spheroids were developed for 5 days in the absence or presence of 50% of MC-conditioned media (CM) or different amounts (10-50%) of PBS. Melanoma cell numbers were determined after trypsinization of the spheroids and staining with trypan blue. Data shown represent mean values ± SEM from a representative experiment. One-way ANOVA, Dunnett’s multiple comparisons test. \**P* < 0.05, \*\**P* < 0.01, \*\*\*\**p* < 0.0001.

## Author Contributions

MG conceived of the study, designed, planned, and performed the experimental work, interpreted data, and contributed to the writing of the manuscript; TN performed experimental work and interpreted data; TH contributed to the design of experiments; HW contributed to the design of experiments; ML performed experimental work; AP contributed to the design of the work, interpreted data and contributed to the writing of the manuscript; GP conceived of the study, planned and interpreted the experiments and wrote the manuscript. All authors read the manuscript and provided input.

## Acknowledgements

We thank the SciLifeLab (Uppsala) for excellent technical support, and Jeremy Adler (BioVis Biological Visualization platform; Uppsala University) for excellent assistance with the sample preparation for the confocal analysis and with subsequent image analysis.

## Availability of Data and Material

The data that support the findings of this study are available within the article. Data and material will be made available on reasonable request to the corresponding authors.

## Abbreviations

MC: mast cell
BMMC: bone marrow-derived mast cell
ECM: extracellular matrix
DCT: L-dopachrome tautomerase

